# Enhancement of vestibular motion discrimination by small stochastic whole-body perturbations in young healthy humans

**DOI:** 10.1101/2022.08.15.504006

**Authors:** Barbara La Scaleia, Francesco Lacquaniti, Myrka Zago

## Abstract

Noisy galvanic vestibular stimulation has been shown to improve vestibular perception in healthy subjects. Here, we sought to obtain similar results using more natural stimuli consisting of small-amplitude motion perturbations of the whole body. Thirty participants were asked to report the perceived direction of antero-posterior sinusoidal motion on a MOOG platform. We compared the baseline perceptual thresholds with those obtained by applying small, stochastic perturbations at different power levels along the antero-posterior axis, symmetrically distributed around a zero-mean. At the population level, we found that the thresholds for all but the highest level of noise were significantly lower than the baseline threshold. At the individual level, the threshold was lower with at least one noise level than the threshold without noise in 87% of participants. Thus, small, stochastic oscillations of the whole body can increase the probability of detecting subthreshold vestibular signals, possibly due to stochastic resonance mechanisms. We suggest that, just as the external noise of the present experiments, also the spontaneous random oscillations of the body associated with standing posture are beneficial by enhancing vestibular thresholds with a mechanism similar to stochastic resonance. The results are also relevant from a clinical perspective, since they raise the possibility of improving motion perception in people with elevated thresholds due to aging or vestibulopathy by means of small-amplitude motion perturbations.

**HIGHLIGHTS:**

- Small-amplitude motion perturbations of the whole body improve vestibular perceptual thresholds of motion discrimination in young healthy people
- Improvements occur at optimal levels of noise amplitude, idiosyncratic to each subject
- The findings are consistent with the phenomenon of stochastic resonance
- The new method can applied to people with elevated thresholds due to aging or vestibulopathy

## INTRODUCTION

By monitoring 3-dimensional angular velocities and gravito-inertial accelerations of the head, the vestibular system contributes to keep clear vision and postural equilibrium (Angelaki and Cullen 2008). Vestibular information is also critical for the perception of head position and displacement, and therefore for our sense of spatial orientation (Merfeld 2012). Thus, discriminating forward from backward direction of passive motion in darkness is a spatial orientation task that heavily relies on vestibular cues. The precision of head motion perception can be quantified by means of the psychometric functions for the discrimination of head-centered passive translations and tilts (Merfeld 2011). The psychometric function yields estimates of the individual vestibular threshold, that is, of the minimum amount of motion necessary to reliably perceive the direction of motion.

In young persons, vestibular thresholds are generally low, denoting great precision of motion discrimination (for a review, see Diaz-Artiles and Karmali 2021). The thresholds progressively increase after the age of about 40 years (Kingma 2005; Roditi and Crane 2012; Bermudez Rey et al. 2016). Since they represent a sensitive measure of vestibular function, they help to identify specific peripheral and central vestibular deficits (e.g., Lewis et al. 2011; Agrawal et al. 2013; Priesol et al. 2014; Bremova et al. 2016; Diaz-Artiles and Karmali 2021; Kobel et al. 2021b). Moreover, there is a correlation between vestibular thresholds and postural stability: higher thresholds tend to be associated with greater postural sway even in young healthy people (Karmali et al. 2021). In elderly people, higher vestibular thresholds correlate with balance test failures (Bermudez Rey et al. 2016), which are known to predict the likelihood of falls in everyday life (Agrawal et al. 2009). Therefore, it would be important to have experimental protocols that can ameliorate the discrimination by lowering the vestibular thresholds, especially in view of clinical applications.

Although external noise usually represents an undesirable disturbance, there exist specific cases of “good” noise improving threshold-like systems. Thus, low amplitude noise added to muscle spindles (Cordo et al 1996), cutaneous receptors (Collins et al. 1996a) or vestibular hair cells (Flores et al. 2016) enhances their responses to weak stimuli. Also, low amplitude noise added to visual (Simonotto et al. 1997), auditory (Jaramillo and Wiesenfeld 1998), tactile (Collins et al. 1996b) stimuli or added directly to cortical networks (Van der Groen and Wenderoth 2016) can improve the sensory thresholds. These observations are often interpreted in the context of stochastic resonance (SR). SR consists in the phenomenon whereby random noise in a nonlinear system enhances detection and transmission of weak signals in the system (Benzi et al. 1982; Gammaitoni et al. 1998). SR predicts that, when using different levels of noise, one obtains a maximum performance at some optimal noise level (Moss et al. 2004; McDonnell and Abbott 2009).

Results compatible with SR have been observed also for head motion discrimination by adding stochastic galvanic vestibular stimulation (GVS) on top of the tested motion stimuli in young healthy participants (Galvan-Garza et al., 2018; Keywan et al., 2018, 2019, 2020a). GVS consists in applying electrical noise to the vestibular system by means of electrodes placed at the mastoids (Fitzpatrick and Day 2004). Although very interesting, these results have been somewhat inconsistent. Thus, GVS significantly decreased roll-tilt thresholds at 0.2 Hz in one study (Galvan-Garza et al., 2018), whereas it did not significantly affect the thresholds at 0.2 Hz but only at 0.5 Hz and 1 Hz in another study (Keywan et al., 2018). Still another study (Keywan et al., 2019) showed that GVS reduced the thresholds for inter-aural translation of upright participants (stimulating the otoliths) but not the thresholds for yaw-rotation with the head pitched forward 71° (primarily resulting in stimulation of the semicircular canals), suggesting that GVS mainly affects otolith-mediated perception (Zink et al. 1998). However, an electrophysiological study in macaques showed that GVS produces robust and parallel activation of both canal and otolith primary afferents, resulting in constant GVS-evoked neuronal detection thresholds across all afferents (Kwan et al. 2019). This study also showed that afferent tuning differs for GVS versus natural motion stimulation, due to the fact that GVS bypasses the biomechanics of both the semicircular canals and the otolith organs (Kwan et al. 2019).

Given that GVS elicits unnatural vestibular afferent inputs, it would be important to be able to enhance head motion discrimination by adding perturbations that resemble natural motion stimuli. A few prior studies tested motion perception using mechanical noise, instead of electrical noise. Wide-spectrum vibrations applied directly to the mastoid did not significantly change the threshold for yaw rotation at 1 Hz (Kabbaligere et al. 2018). Vertical whole-body oscillations (6 Hz; 1-2 mm amplitude; peak acceleration of about 140 cm/s2) resulted in horizontal heading-direction thresholds higher than without the perturbation (Rodriguez and Crane 2018). A few studies tested whether a prolonged exposure to conditioning passive motions modifies perceptual thresholds in a subsequent session. Thus, 10-minutes of conditioning stimuli consisting of high-amplitude stochastic yaw rotations (0.5–2.5 Hz; up to 300 deg/s^2^) increased substantially yaw perceptual thresholds (Fitzpatrick and Watson 2015). By contrast, 20-minutes of conditioning stimuli consisting of small-amplitude, subliminal interaural translations (1 Hz sinusoid; 2 cm/s^2^) significantly reduced the thresholds for motion discrimination along the same axis (Keywan et al. 2020b), as well as along the naso-occipital axis (Keywan et al. 2022).

Here, we re-examined the issue of the potential effects of small-amplitude motion perturbations applied during the motion discrimination task. The experiment involved a direction-recognition task, in which the subject reported the perceived direction of the motion (two-alternative forced-choice). First, we determined the individual perceptual thresholds to antero-posterior translations in a baseline condition. Next, we carried out 5 blocks of trials where we measured again the perceptual thresholds to these translations while simultaneously applying small, stochastic perturbations along the antero-posterior axis, symmetrically distributed around a zero-mean. The power of the perturbations was proportional to the power of the acceleration signal reliably perceived by the participant in the baseline condition (vestibular threshold). The proportionality coefficient was set at 5 different values, including a zero-level to verify the consistency of the baseline threshold estimate. We hypothesized that the probability of detecting a subthreshold signal was higher in the presence of noise at an optimal level than in the absence of noise, possibly due to SR effects. As a result, the motion discrimination thresholds should be lower with noise than without.

## MATERIALS AND METHODS

### Participants

Thirty subjects (21 females; 9 males; 24.6 ± 7.4 years, mean ± SD) participated in the study. They gave written informed consent to procedures approved by the Institutional Review Board of Santa Lucia Foundation (protocol n. CE/PROG.757), in conformity with the Declaration of Helsinki (World Medical Association, 2013) regarding the use of human participants in research. All participants had normal or corrected-to-normal vision, no history of psychiatric, neurological or vestibular symptoms, dizziness or vertigo, motion-sickness susceptibility, major health problems or medications potentially affecting vestibular function. Sample size was calculated to detect an effect size of 0.8 (Cohen’s d, estimated from Keywan et al., 2019 and pilot data with the current setup), by considering paired t-test (R package pwr) with a power of 0.8 and an alpha of 0.05/5 (multiple testing correction), allowing for 25% loss of participants due to various reasons.

### Setup

Participants sat in an upright position in a padded racing chair mounted on top of a 6DOF hexapod motion platform (MOOG MB-E-6DOF/12/1000Kg, East Aurora, New York, USA). A 4-point harness held their trunk securely in place. A medium density foam pad under their feet minimized plantar cues about body displacement. Their head was positioned against a headrest in a comfortable posture, centered left to right relative to the earth-vertical and up to down relative to the antero-posterior direction of translation using external landmarks. It was then held in place by means of a tight headband. We monitored 3D position and orientation of both the platform and the participant’s head at 200 Hz by means of the Optotrak 3020 system (Northern Digital, Waterloo, Ontario). To this end, 4 non-coplanar infrared emitting markers were attached to the right side of both the chair and the participant’s head. In the latter case, 2 markers were attached roughly in correspondence of the zygomatic bone, one marker was attached on the tragus, and one marker over the parotid gland. All markers were visible in the initial calibration phase of the experiment. During the test phase, the participants wore active noise-canceling headphones (Bose Noise Cancelling Headphones 700) to mask the acoustic noise from the motion platform. Since the headphones obscured the tragus marker, its virtual position was estimated from a rigid body model of the head constructed from the calibration data. Through the headphones, we also delivered task instructions. To eliminate visual cues during the test phase, the participants kept their eyes closed in the light-tight room. They entered the responses via two buttons of a wireless gamepad. Participants always wore face masks according to the Institution regulations related to COVID-19.

### Motion stimuli

Stimuli differed in the baseline trials and in the trials with non-zero perturbations (see Fig. 1 and *Protocol* below). In the former case, the stimuli were pure single cycles of sinusoidal acceleration along the antero-posterior axis (roughly corresponding to the naso-occipital axis), parallel to the earth horizontal plane, in either forward or backward direction:

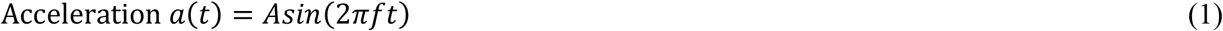

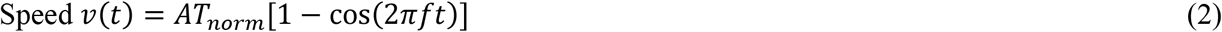

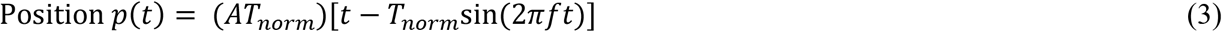

where *A* is the acceleration amplitude with frequency f = 1 Hz, *T*_*norm*_ =*T/*(2π), and *T* = 1 s is the duration of the motion cycle. In each trial, the value of *A* was adjusted based on an adaptive staircase (see *Procedure*). In the following, the reported thresholds correspond to the peak speeds of the smallest stimulus that was reliably perceived by the participants. Because acceleration, speed, and position are all proportional between each other in Eqs. 1-3, the results are unchanged if presented as displacement thresholds, velocity thresholds, or acceleration thresholds (Chaudhuri et al. 2013).

**Figure 1.**
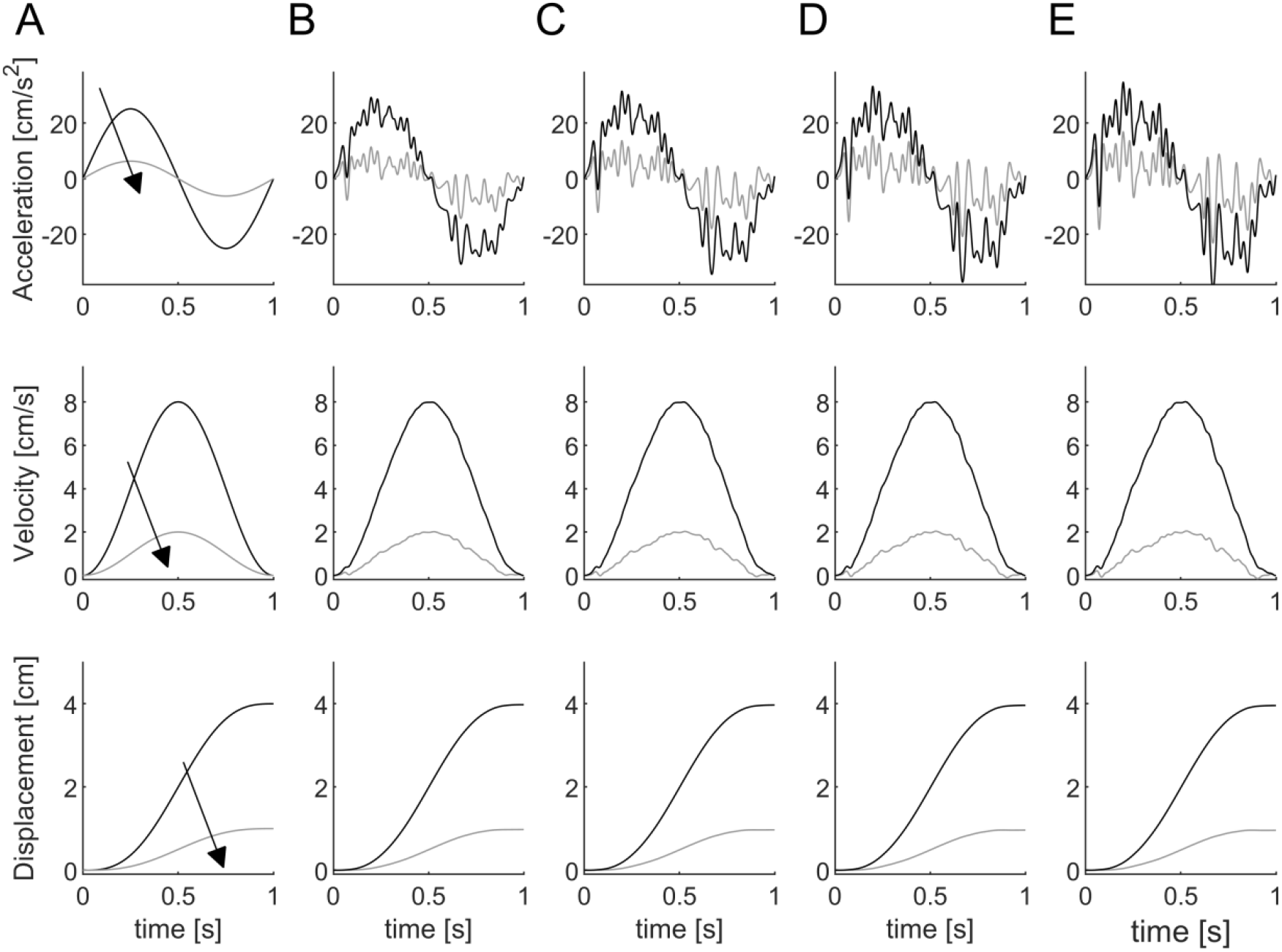
Motion stimuli consisting of a single cycle of 1 Hz sinusoidal accelerations along the antero-posterior direction (top row), the corresponding velocities (middle row) and displacements (bottom row). A: Black and gray correspond to unperturbed stimuli with peak velocities of 8 and 2 cm/s, respectively. The arrow indicates the direction of changes following a theoretical adaptive staircase. B: Perturbed stimuli with peak velocities of 8 and 2 cm/s in the noise condition with intensity proportional to a vestibular threshold of 2 cm/s and *k*=0.5. C, D and E: The same of panel B but *k*=1, 1.5 and 2, respectively.

In the trials with perturbations, noise of different intensity was superimposed onto the same sinusoidal test stimuli used in the baseline trials (Fig. 1). Noise consisted of random fluctuations of acceleration along the antero-posterior axis (the same as that of the test stimuli). Noise power was proportional to the power of the acceleration stimulus corresponding to the individual threshold determined for each participant during the baseline condition. Specifically, we generated offline (at 1 kHz) white noise *ε(t)*, 1-s duration, which was band-pass filtered within 1.8 Hz-30 Hz (infinite impulse response filter of order 20, Matlab function *bandpassiir*), reduced to zero-mean by subtracting the mean value, and normalized to unit-power. This signal was scaled in amplitude as a function of the variance *δ*_*2*_ of the acceleration at the individual baseline threshold:

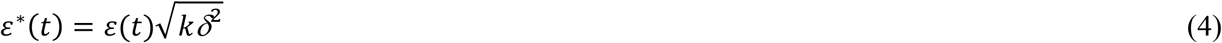

The proportionality constant *k* was 0, 0.5, 1, 1.5, or 2 in different blocks of trials. The condition with *k* = 0 was identical to the baseline, and represented an experimental control to verify the consistency of the baseline threshold estimate. The overall acceleration stimuli applied in the trials with noise were:

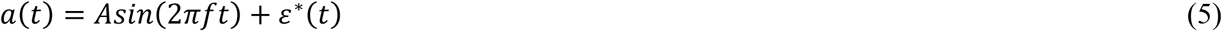

Notice that the value of *A* changed from trial to trial according to the adaptive staircase in the same manner in the trials with and without noise, but the added noise *ε*^∗^(*t*) was the same in all trials of a given block. However, as noticed above, the noise intensities were idiosyncratic to each participant, depending on the individual baseline thresholds.

By design, the noise *ε*^∗^(*t*) was symmetrically distributed around a zero-mean not to provide any directional cue independently of the potential interaction with subthreshold test stimuli.

All position profiles were programmed in LabVIEW 2020 (National Instruments, Austin, Texas, USA) with custom-written software, and input to the MOOG controller at 1 kHz.

### Procedure

To determine the perceptual thresholds, we used a 3Down-1Up adaptive staircase (Leek 2001; Grabherr et al. 2008; Karmali et al. 2016). Initial peak speed of the unperturbed motion stimulus was 8 cm/s, above the presumptive threshold in each participant (Diaz-Artiles and Karmali 2021). The peak acceleration was 25.13 cm/s^2^, and the maximum displacement was 4 cm. Until the first mistake, the stimulus was halved after three correct responses at each level. From this point onward, the size of the change in stimulus amplitude was determined using parameter estimation by sequential testing (PEST) rules (Taylor and Creelman 1967). The minimum step size was 0.38 dB (i.e., 1.25 log_10_2), and the maximum step size to 6.02 dB (i.e., 20 log_10_2). Motion direction (forward or backward) was randomized in each trial. The randomization procedure ensured that there was the same number of forward and backward motions every 20 trials. After each stimulus, participants indicated the perceived direction of motion (forward or backward, two-alternative forced-choice direction recognition task). When they were unsure of the direction, they were asked to make their best guess. No feedback was provided as to the correctness of the responses. In each trial starting from the 25th, we iteratively fitted a running psychometric curve with a generalized linear model (GLM) and we computed the coefficient of variation of the σ parameter (see *Data analysis*). The block of trials was terminated after 100 trials or when σ reached 0.2, whichever occurred first.

### Protocol

Participants received detailed instructions about the procedure prior to the experiment. However, neither the possible presence of noise perturbations nor the purpose of the experiment was disclosed. All participants performed 6 blocks of trials, split in 2 sessions on 2 separate days (separated by about 8 days) to avoid fatigue, with 3 blocks of trials in each session. Before each session, head position and orientation were recorded over 5 s during a calibration phase and the average values used as a reference for the following trials. Calibration was repeated during the session if necessary (e.g., when the participant stepped down the chair to rest). Next, six suprathreshold (8 cm/s speed, 1 Hz), practice trials without noise were administered to make sure that the participant was comfortable with the task. On day 1, the first block always involved the determination of the baseline threshold using motion stimuli without noise perturbations (Eq. 1). This baseline threshold was used to compute the individual noise level for the trials with added perturbations. The next 5 blocks of trials involved the 5 noise levels (*k* = 0, 0.5, 1, 1.5, 2 in Eq. 4) in randomized order, counterbalanced across participants. The condition with *k* = 0 was the same as the baseline, and it was randomly interspersed with the others with non-zero noise. Each block of trials lasted about 13 minutes, with about 10-minutes rest breaks between blocks. No participants -interviewed after the end of the experiment on day 2-reported having being aware of the presence of noise perturbations.

The time sequence of events during each trial was the following. The participant pressed a button when ready for the trial after hearing a pre-recorded voice message. Before starting the motion stimuli, we checked that the head had not moved appreciably relative to the reference of the calibration phase. To this end, we acquired 3D head position and orientation over 500 ms, and computed the mean shift relative to the reference. The shift in position was calculated as the 3D distance of the tragus from its reference position. The tilt was calculated separately for roll, pitch and yaw from the reference angles. If the shift was less than ≈ cm for position and ≈ 5 ° for tilt, platform motion could start. Else, the participant’s head was repositioned within the described tolerance window, and the trial started again with the voice message. The time interval between the button press and the motion start was 2 s. Two consecutive sounds (each with frequency of 500 Hz, duration of 125 ms) signaled the end of platform motion. Within the next 2 s epoch, participants had to indicate the perceived motion direction by pressing the fore or aft button of the hand-held gamepad, depending on whether they perceived a forward or backward motion, respectively. If they responded too early or too late, a different sound (frequency 250 Hz, duration 800 ms) signaled the error, and the response was considered incorrect. They were maintained in the final position for 3 s, then they were moved back to the initial position with the same kinematics of the last stimulus but in the opposite direction. Thus, the time interval between the start of platform motion and the return to the initial position was 5 s. The minimum inter-trial interval was 3.34 s, ensuring vestibular wash-out. During the platform motion, a running average of 3D head position and orientation over 50-ms consecutive intervals was computed on-line. If the head shifted by >0.5 cm or rotated by >2.5° (in either roll, pitch or yaw) relative to the chair (and platform) over any 50-ms interval, the trial was discarded and repeated.

### Data analysis

Data analyses were performed with Matlab 2021b (The MathWorks, MA, USA). We fit a Gaussian cumulative distribution psychometric function with standard deviation (σ) and mean (μ) to the responses with a maximum likelihood estimate via a GLM and a probit link function (Merfeld, 2011). The threshold parameter represents the “one-sigma” vestibular threshold and corresponds to (1) the standard deviation of the underlying distribution function and (2) the stimulus level that would be expected to yield 84% correct performance in the absence of bias (Merfeld 2011). Each data set was also fit using a bias-reduced generalized linear model (BRGLM, Matlab function *brglmfit*) to correct potential misestimates of σ when fitting serially dependent data (Chaudhuri and Merfeld 2013; Karmali et al. 2016). Moreover, we fit the psychometric function using a lapse-identification algorithm (LIA, deltadeviance method, Clark and Merfeld 2021) based on a standard delete-one jackknife procedure to identify probable spurious datapoints of the staircase (Tukey 1958). Lapses are errors made by participants independently of the test stimuli, such as those due to inattention, fatigue etc.

### Characterization of mechanical stimuli

We carried out separate tests to verify that the mechanical perturbations were equivalent when applied during forward and backward translations, so as not to provide any directional cue independently of the potential interaction with subthreshold test stimuli. To this end, 3D linear accelerations were recorded at 200 Hz with an MPU-6050 sensor (TDK InvenSense, San Jose, California, USA, operated at full scale range of ±2g) attached to the base of the MOOG platform under the chair, while position of the markers on the chair was recorded by the Optotrak at the same rate. Data were acquired during conditions replicating those experienced by our participants when they were close to a typical value of perceptual threshold (see *Results*). Thus, we applied 1-Hz single-cycle sinusoidal accelerations with peak velocity of 2 cm/s in either forward or backward direction, and noise intensity proportional to the vestibular threshold of 2 cm/s and *k*=1 (Eq. 4). In each trial, we recorded a 2 s time epoch extending 0.5 s before and after the motion trajectory. We performed 100 trials for each motion direction (forward and backward). Data analysis followed that of Chaudhuri et al. (2013) for both positions and accelerations. To minimize potential biases and drifts, the mean value for the first and the last 0.2 s was subtracted from the data of each trial. Then, we computed the average for each of the three orthogonal linear acceleration (position) components at each instant in time over all 100 trials for each direction. Using the three average components of acceleration (position), we computed the module of acceleration (displacement) in forward and backward direction at each instant in time. We found that the module of acceleration and position were not significantly different between forward and backward direction. Thus, the mean difference between the module of acceleration in forward and backward direction was 0.06 cm/s^2^ [95%CI -0.04 0.17], while the mean difference between the module of displacement in forward and backward direction was 0.0002 cm [95%CI -0.0001 0.0004].

We also used the results of these tests to verify that the actual position profiles generated by our MOOG platform matched the programmed profiles, since mechanical dynamics might result in a mismatch (Karmali et al. 2014). We found that the mean difference between the module of position measured by the Optotrak and the module of the position signal input to the MOOG was -0.0010 cm [95%CI -0.0853 0.0833] and -0.0008 cm [95%CI -0.0853 0.0836] in forward and backward direction, respectively.

### Statistics

Statistical analyses were performed in R (4.0.2). Individual threshold values were first log transformed, because the results with GLM and BRGLM demonstrated a lognormal distribution in all participants, consistent with previous reports (Grabherr et al., 2008). We used the Shapiro–Wilk test to verify the normality of distribution of data. We also verified whether there were outlier participants by applying the R function identify_outliers (rstatix package) to the individual threshold values of the baseline block.

To test for effects of noise on motion discrimination, log-transformed thresholds were subjected to repeated-measures analysis of variance (RM-ANOVA) with noise intensity as the within-subjects factor (six levels, baseline and *k* = 0, 0.5, 1, 1.5, and 2). Effect size was computed as partial eta square 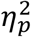. Additionally, preplanned comparisons testing performance for each of the 5 noise intensities (*k* = 0, 0.5, 1, 1.5, or 2) against the baseline were performed and corrected for multiple comparisons by means of the Bonferroni method (using the R function pairwise_t_test).When the log-transformed thresholds were not normally distributed (with the lapse-identification LIA), we used non-parametric statistics to test the effects of noise (Friedman test). Post-hoc corrections for multiple comparisons were performed with Wilcoxon test (R function wilcox_test).

Population responses were also analysed by means of a Generalized Linear Mixed Model (GLMM) that separately accounts for the random effects due to inter-subject variability and the fixed effects due to the experimental variables (Moscatelli et al., 2012). To this end, we fitted the data of the different participants and experimental conditions with the following GLMM:

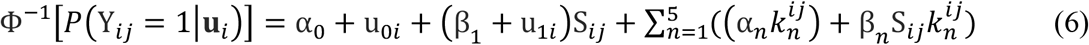

The left side of the equation is the probability that participant *i* in trial *j* reported that the platform motion was in the forward direction, with Ф^−1^ being the probit transform of this probability (i.e., the inverse of the cumulative Gaussian function). The right side of the equation is a linear combination of the fixed (α and β) and random (u) effects predictors. Specifically, *S*_*ij*_ is the amplitude of the stimulus and *k*_*n*^*ij*^_ (with *n*=1:5) are the categorical predictors coding for the perturbed conditions (with noise level of *k*=0, 0.5, 1, 1.5, 2). The first unperturbed condition represented the baseline in the model. The fixed effects estimated the effect of the experimental variables common to all participants. The fixed effects α_0_, and β_0_ correspond to the intercept and the slope of the baseline condition. The inverse of the slope represents the threshold (σ): the higher the slope, the lower the threshold. The slope of the condition with a given level of noise (k(n)) is equal to the sum of β_0_ and β_n_. If the threshold of a condition with noise (σ_n_) was not significantly different from the baseline (σ_0_), then β_n_ would not be significantly different from zero (the null hypothesis). The fixed-effect parameters α_n_ provided an adjustment to the intercept in each noise condition. The random-effect parameters u_0i_ and u_1i_ estimated the heterogeneity between participants.

Alpha was set to 0.05 for all statistics.

## RESULTS

Each participant underwent 6 experimental conditions with 2 blocks of trials involving unperturbed stimuli (the baseline and the control with *k*= 0) and 4 blocks of trials involving perturbed motion stimuli (*k*=0.5, 1, 1.5, 2). While the sinusoidal motion stimuli were the same for all participants in all trials (both unperturbed and perturbed), the noise intensities in the perturbed trials were idiosyncratic to each participant, depending on the individual baseline thresholds (see *Methods*).

On average, in each block participants performed 77.5 valid trials (11.6 SD, n=180 [30 participants x 6 blocks)]) before reaching the preset target. Due to inattention or head movements outside the tolerance window (see *Methods*), there was also an average of 4.03 (6.2 SD, n=180) invalid trials that were discarded and repeated during the block. The number of both valid and invalid trials did not depend significantly on the 6 experimental conditions (P=0.5 and 0.23 respectively, Friedman test).

At the start of the trial, the mean shift of the head in 3D relative to the calibration reference was 0.3 cm (SD=0.4 cm, n=180), and -0.51°, 0.89°, 0.57° (SD 1.36, 2.76, 1.48, n=180) in roll, pitch and yaw, respectively. During the platform motion, the mean shift of the head in 3D relative to the platform was 0.14 cm (SD 0.04 cm, n=180), and 0.96°, 0.43°, 1.43° (SD 0.29, 0.22, 0.38, n=180) in roll, pitch and yaw, respectively. Neither the head shift at trial start nor that during platform motion depended significantly on the experimental condition (all P>0.13, Friedman test).

The mean amplitude of the sinusoidal test stimuli (i.e. the platform displacement) in correspondence of the baseline threshold was 1.14 cm (SD=0.34, n=30 participants). By comparison, the mean amplitude (maximum – minimum) of the noise perturbations was 0.027 cm (SD= 0.008, n=30), 0.038 cm (SD= 0.011), 0.047 cm (SD= 0.014), 0.054 cm (SD= 0.016) for noise level *k*=0.5, 1, 1.5, 2, respectively. Thus, the noise amplitude was less than 5% of the smallest stimuli that were reliably perceived by the participants.

### Thresholds

We found that the log-transformed thresholds computed with GLM were normally distributed for all 6 experimental conditions (baseline and *k* = 0, 0.5, 1, 1.5, and 2, all P-values > 0.178, Shapiro–Wilk test). The thresholds of each participant and each condition are plotted in Fig. 2. There was considerable inter-subject variability, and there were no outlier participants for the baseline condition. Importantly, the threshold in 26/30 participants was lower with at least one noise intensity than the corresponding values in both unperturbed conditions (baseline and control, grey shading in Fig. 2), indicating that low amplitude noise added to vestibular stimulation can improve the perception of motion direction.

**Figure 2.**
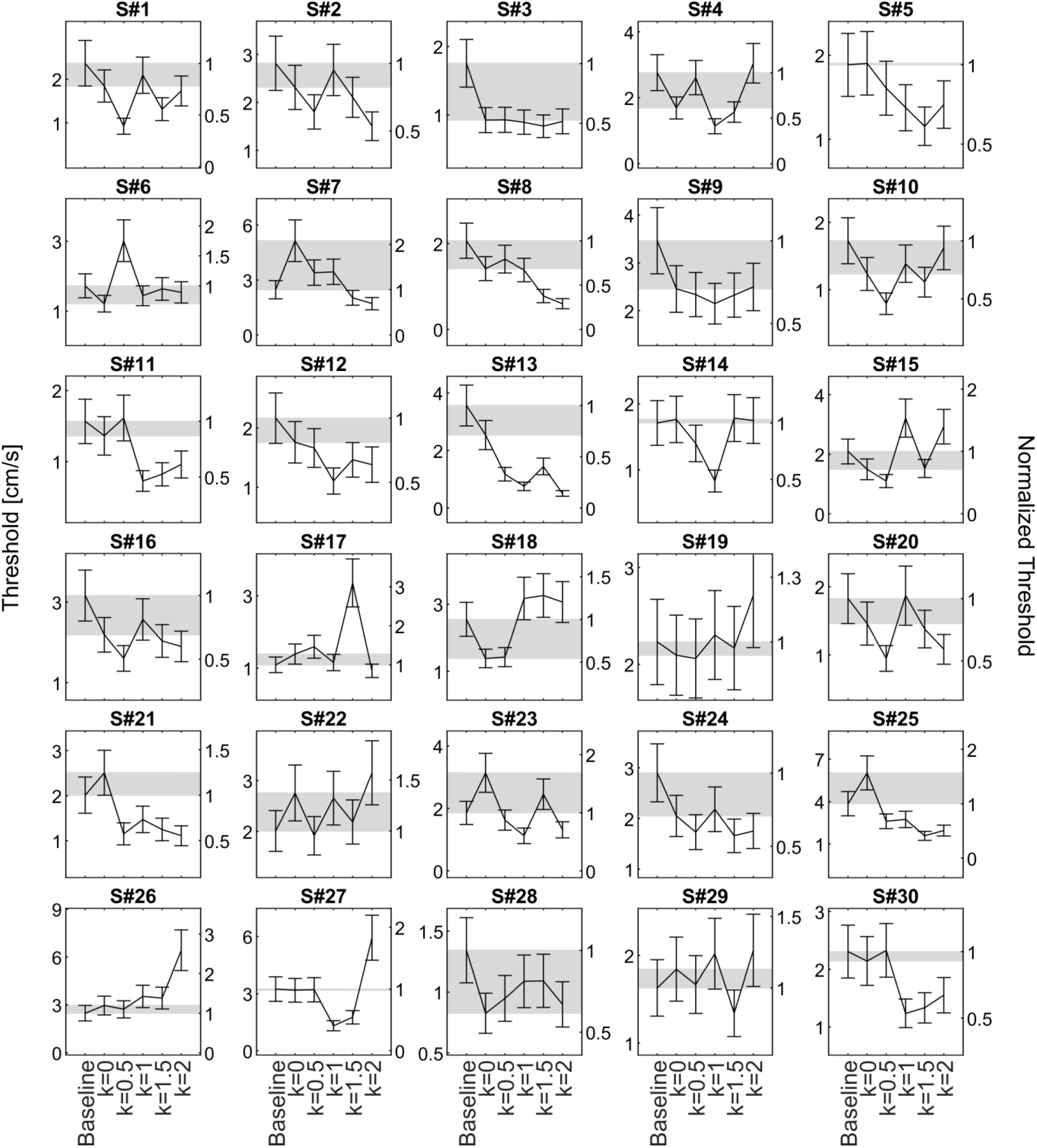
Motion discrimination thresholds of each participant in all experimental conditions (error bars indicate the standard deviation of the threshold estimate). Grey shading indicates the range of threshold values in unperturbed conditions (baseline and control with *k*=0). Left axis indicates the absolute value of threshold, the right axis indicates the threshold value normalized by the baseline threshold.

A RM-ANOVA over the responses of all 30 participants showed that the threshold was significantly different as a function of the 6 experimental conditions (F(5,145) = 4.17, P = 0.001, 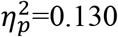). Post-hoc tests showed that neither the threshold for the control condition (*k*= 0) nor that for the highest level of noise (*k*= 2) differed significantly from the baseline threshold (uncorrected P=0.081 and P =0.013, respectively). By contrast, the thresholds with noise levels *k*= 0.5, 1 and 1.5 were all significantly lower than the baseline (all P < 0.009 after Bonferroni correction, Fig. 3, Table 1).Thresholds were also computed with the Generalized Linear Mixed Model (GLMM) that separately accounts for the random effects due to inter-subject variability and the fixed effects due to the experimental variables (Eq. 6). The GLMM confirmed the previous results by showing that the slope of the responses (the inverse of the slope corresponds to the threshold, see *Methods*) was significantly higher (implying a lower threshold) for the conditions with noise levels *k*= 0.5, 1 and 1.5 than the slope for the baseline (P < 0.001). By contrast, the slopes of the responses for the control condition (*k* = 0) and for the highest level of noise (*k* = 2) were not significantly different from the slope of the baseline (all P>0.09).

**Table 1.**
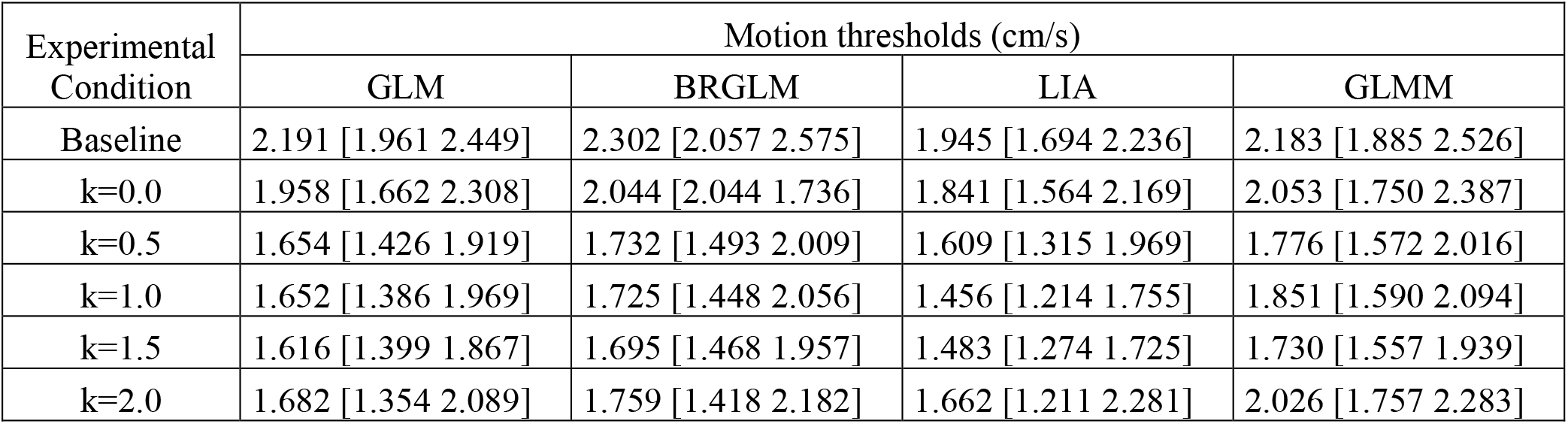
Vestibular thresholds computed with different methods (see text). For GLM, BRGLM and GLMM, the values correspond to the geometric means. For LIA, the values correspond to the medians, since the data were not normally distributed. 95% confidence intervals are between brackets.

**Figure 3.**
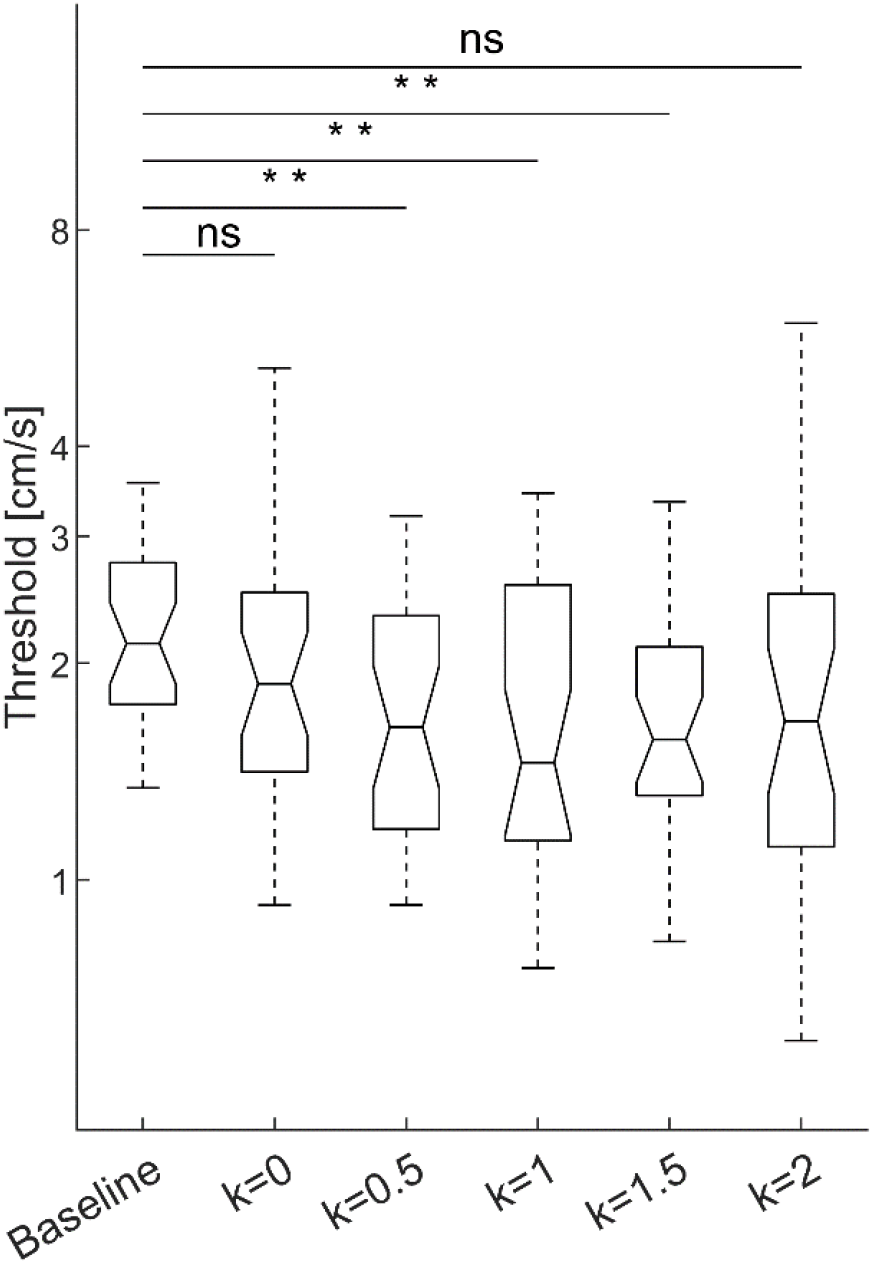
Effects of different levels of noise on motion thresholds at population level. The thresholds with noise intensity *k*= 0.5, 1 and 1.5 were significantly lower than the baseline threshold (**, P<0.01). The threshold for the control condition (*k*= 0) and that for the highest level of noise (*k*= 2) were not significantly different from the baseline threshold. In the box-and-whisker plots, the box corresponds to the median and the 25th and 75th quartiles, and the whiskers show the 5th and 95th percentile.

We obtained very similar results by calculating the thresholds with the bias-reduced generalized linear (BRGLM) model or with the lapse-identification algorithm (LIA, Table 1). In both cases, the threshold was significantly different as a function of noise (with BRGLM, F(5,145) = 4.281, P = 0.001,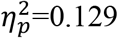 with LIA, Friedman test χ^2^(5) = 13.1, n=30, P = 0.023, 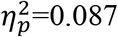).

As it is evident from Fig. 2, there was a high inter-individual variability in the optimal noise level, that is, the noise level leading to the best detection performance. Moreover, as we remarked before, the noise intensities in the perturbed trials were idiosyncratic to each participant, depending on the individual baseline thresholds. Therefore, we compared the lowest thresholds obtained in each individual (irrespective of the noise level, *k*= 0.5, 1, 1.5 or 2) with the thresholds obtained in the unperturbed conditions (baseline and *k*=0, Fig. 4). RM-ANOVA showed that the threshold was significantly lower with the optimal individual noise than without noise (F(2,58) = 42.49, P<0.0001, 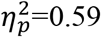; post-hoc tests all P <0.0001 after Bonferroni correction). On average, the optimal noise reduced thresholds by 42% relative to the thresholds without noise.

**Figure 4.**
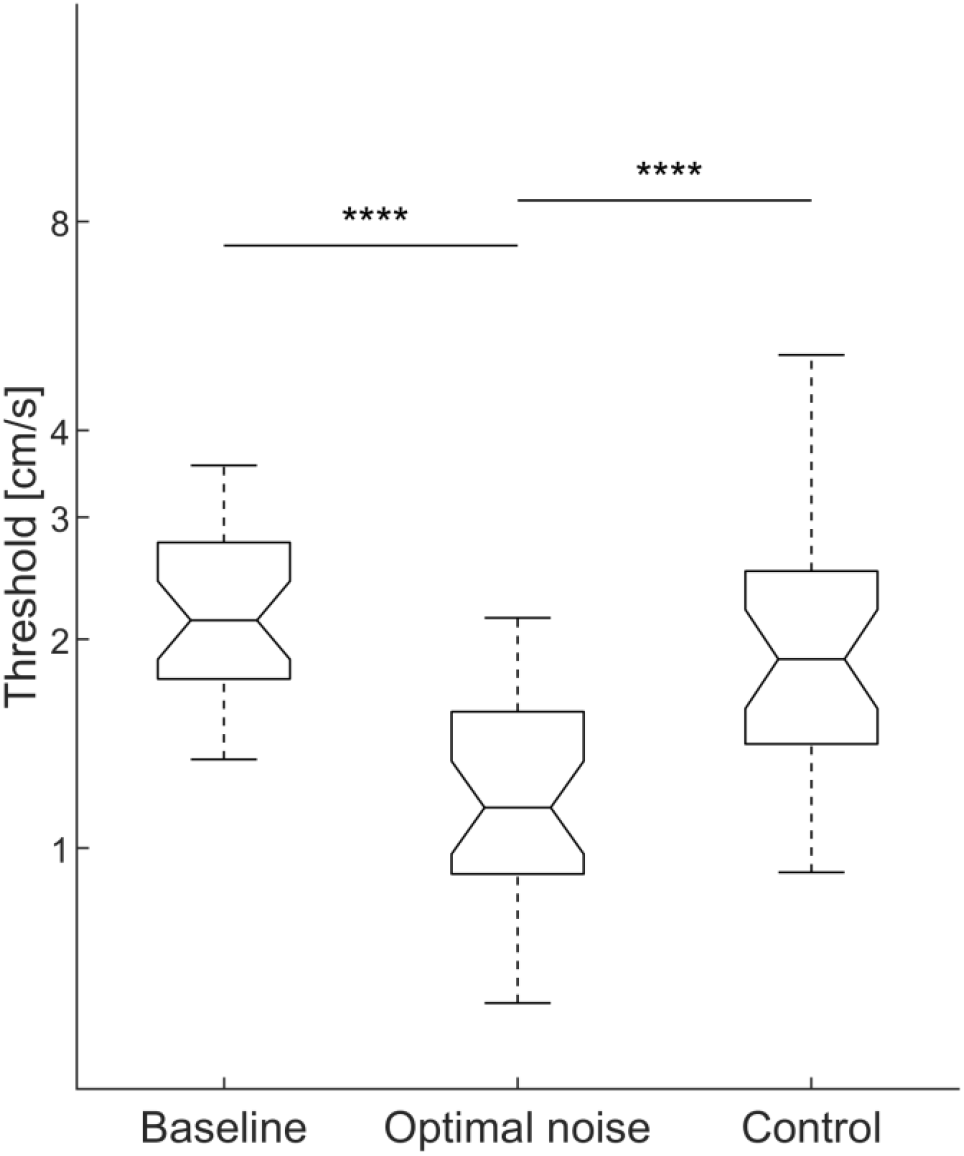
Comparison of population thresholds at optimal noise level with the thresholds in the unperturbed conditions (baseline and control). Optimal results were obtained by pooling together the lowest thresholds obtained in each participant irrespective of the noise level at which they were observed. The differences relative to the unperturbed conditions were statistically significant (****, P<0.0001). Same format of plot as in Fig. 3.

## DISCUSSION

We applied small-amplitude motion perturbations during a motion discrimination task at different levels of noise. At the population level (30 participants), we found that the thresholds for all but the highest level of noise were significantly lower than the baseline threshold. At the individual level, the threshold was lower with at least one noise level than the threshold without noise in 26/30 (87%) participants. We suggest that stochastic mechanical oscillations of the whole body can increase the probability of detecting subthreshold vestibular signals, possibly due to stochastic resonance (SR) effects.

### Threshold values

With the noise conditions, we randomly interspersed a control with zero noise to verify the consistency of the baseline threshold estimate. At the population level, we found no statistically significant difference of threshold between the control and the baseline. This result is consistent with previous reports showing limited session-to-session, intra-subject variability of the motion discrimination thresholds (Hartmann et al. 2013; Clark et al., 2018). On the other hand, we found large inter-subject variability of the thresholds both in the unperturbed conditions (baseline and control) and in the noise conditions. Inter-subject variability of vestibular thresholds has also been reported in previous studies (Bermudez Rey et al. 2016; Diaz-Artiles and Karmali 2021).

As for the absolute values of the motion thresholds, these may not be easily comparable across different studies because of methodological and individual differences (Diaz-Artiles and Karmali 2021). The motion thresholds we found roughly overlap those reported by Bremova et al. (2016) for similar stimuli (1 Hz forward-backward translations) and setup. However, our thresholds were higher than those reported by Kobel et al (2021a). Since we used similar motion waveforms in the unperturbed conditions and similar procedures to compute the thresholds as Kobel et al (2021a), the differences presumably depend on either the setup or subjective factors.

### Stochastic resonance

The present results are compatible with the presence of stochastic resonance (SR). In SR, the addition of a random interference (noise) enhances the detection of weak stimuli or the information content of the signal (Moss et al. 2004). An optimal amount of added noise results in the maximum enhancement, and further increases in the noise intensity do not lead to any enhancement or even degrade detectability. Consistent with this prediction, we found that the threshold at the highest level of noise was not significantly different from the baseline. However, the optimal level of noise differed considerably across participants. In part, this variability might depend on the fact that the specific intensities of the noise were idiosyncratic to each participant, being determined on the basis of the individual baseline thresholds which were quite variable, as noticed above. In part, the variability can be reconciled with the high inter-individual variability in optimal noise levels that has been reported in several previous SR studies (e.g.,Collins et al. 1996; Martínez et al., 2007; Van der Groen and Wenderoth 2016). When we pooled together the lowest thresholds obtained in each participant irrespective of the noise level at which they were observed, we found a reduction of the threshold relative to the conditions without noise (both the baseline and the control) that was statistically robust and strong in magnitude (Fig. 4).

### Comparison with previous studies

Our results confirm previous results obtained by different groups using stochastic galvanic vestibular stimulation (Galvan-Garza et al., 2018; Keywan et al., 2018, 2019, 2020a), and extend the conclusions to more natural vestibular stimuli. These previous studies found significantly lower vestibular motion thresholds with noise than without, and the trends with noise intensity were compatible with SR effects. Similarly to our results, improvements in vestibular thresholds were found within the subject pool, but they were not consistent from subject to subject. Also, the optimal stimulus intensity yielding the largest reduction of threshold could differ substantially across subjects, as was the case in the present study.

The present results differ from those obtained by Rodriguez and Crane (2018). They compared the thresholds for motion discrimination in the antero-posterior direction in the presence or absence of vertical whole-body, quasi-sinusoidal oscillations. They found that the thresholds with the perturbations were systematically higher than without them. However, their perturbations had a different waveform (sinewave) and direction (vertical) than our perturbations (random noise in the antero-posterior direction). More importantly, their perturbations involved an acceleration variance more than 10 times greater than that of our perturbations. As argued by the authors, their perturbations were probably out of range to elicit SR effects.

Our results also differ from those obtained by Chaudhuri et al. (2013). They did not apply external perturbations to the participants, but exploited the intrinsic vibrations of two different motion platforms, a moderate-vibration and a low-vibration system. They did not find significant differences in the motion discrimination thresholds for yaw rotations between the two setups, and concluded that these thresholds are relatively unaffected by vibration when moderate vibration cues are present. Their results are not directly comparable to ours, since they were obtained for a different motion axis (yaw rotation), different stimuli, and different setups (their MOOG is a different model from ours).

Kabbaligere et al. (2018) applied wide-spectrum (1-500 Hz) random vibrations to bilateral mastoids, which are known to stimulate the vestibular system (Young et al. 1977). They did not find significant differences in the threshold for yaw rotation with or without vibration.

Overall, the bulk of the studies indicate that external noise can increase, decrease or leave unaffected the motion recognition thresholds depending on the type and intensity of noise. On the one hand, the conclusions derived from one study may not apply to other studies employing different kinds of perturbations or different motion platforms. On the other hand, the variability of the effects as a function of the noise characteristics is a defining feature of stochastic resonance phenomena (Gammaitoni et al. 1998; Moss et al. 2004; McDonnell and Abbott 2009), and more generally of neural processing of noisy signals (Faisal et al. 2008).

### Putative substrates

We tested motion discrimination in the antero-posterior direction. While this direction roughly corresponded to the naso-occipital axis, we did not check the exact alignment with this axis. Moreover, we measured some (small) variability in head orientation across experiments, which was expected since the head was not rigidly fixed relative to the platform.

Translations along the naso-occipital axis mainly assess utricular function, but also engage the saccules to some extent (Kobel et al. 2021a). Studies have demonstrated that vestibular cues predominate for the perception of motion direction in blindfolded subjects, proprioceptive and tactile cues from the motion platform being absent or minimal (Chaudhuri et al. 2013; Valko et al. 2012). On the other hand, the small-amplitude noise we added to the translations generated vibrations that presumably affected multiple vestibular and non-vestibular (e.g., skin, muscle, visceral) receptors. This noise combined with the small mechanical vibrations intrinsic to the moving Moog platform (Roditi and Crane 2012; Chaudhuri et al. 2013). While humans are extremely sensitive to minute vibrations (Parsons and Griffin 1988), it has been argued convincingly that, in line of principle, vibrations can contribute to motion detection but not to motion direction discrimination tasks (Merfeld 2011; Chaudhuri et al. 2013). Here, given the symmetrical nature of the applied noise, subjects could not extract directional information from the noise *per se*. However, our results indicate that the energy of the noise added to that of subthreshold motion stimuli, thus lowering the thresholds of direction discrimination.

Otolith receptors in the maculae are extremely sensitive, being able to detect displacements of the cilia at atomic scales and correspondingly small accelerations (Howard and Hudspeth 1988). In the maculae, the striola band consists of mainly type I receptors whose hair bundles are weakly tethered to the overlying otolithic membrane. The afferent neurons have irregular resting discharge and have low thresholds to high frequency bone-conducted vibration (Young et al. 1977; Curthoys et al. 2017). We surmise that irregular afferents may mediate SR-like effects due to small vibrations. In a similar vein, it has been suggested that Brownian motion of the hair bundle of the inner hair cells of the cochlea serves to enhance the sensitivity of mechanoelectrical transduction (Jaramillo and Wiesenfeld 1998).

### Limitations

As a first attempt to quantify the effects of small perturbations on the motion direction discrimination, our study has several limitations that could be remedied in future studies. First, because of the constraints on overall testing time (due to COVID-19 regulations), we only investigated the effects of noise on vestibular motion discrimination in the antero-posterior direction. Moreover, we applied noise in the same direction as the motion stimuli to be discriminated. We do not know whether perceptual enhancements would still be obtained with noise applied in a direction different from that of the motion stimuli. For instance, prolonged conditioning stimuli consisting of subliminal interaural translations reduced the thresholds not only for motion discrimination in the same direction but also in the naso-occipital direction (Keywan et al. 2020b; Keywan et al. 2022). However, these conditioning stimuli did not affect significantly the thresholds for yaw rotations (Keywan et al. 2020b), indicating some selectivity for the otoliths.

We only tested one frequency (1 Hz) of motion stimuli to be discriminated. This frequency was chosen based on previous studies showing that discrimination performance is adequate at that frequency (Agrawal et al. 2013; Bremova et al. 2016; Kobel et al 2021a). The frequency range of the applied noise (1.8-30 Hz) was chosen to mimic that of previous studies showing SR-effects on motion discrimination with stochastic galvanic vestibular stimulation (Galvan-Garza et al., 2018; Putman et al. 2021). In the future, one may want to test the effects of mechanical noise in the frequency range of natural head motions during daily locomotion (about 0.5–7 Hz, Carriot et al. 2014).

We only used 4 levels of non-zero noise, again due to the constraints in testing time. The ability to detect a clear-cut U-shaped function as a function of applied noise (the hallmark of SR) strongly depends on the application of several noise levels (Moss et al. 2004; McDonnell and Abbott 2009).

## Conclusions and perspectives

The evidence we provided that small-amplitude motion perturbations can enhance the perceptual discrimination of motion direction has potential implications for both physiology and medicine. From a physiological standpoint, our results suggest the possibility that, just as the external noise that we applied, also the spontaneous random oscillations of the body associated with posture (Duarte and Zatsiorsky 2000) are beneficial by enhancing vestibular thresholds with a mechanism similar to SR. From a clinical standpoint, there is the possibility that the application of small-amplitude motion perturbations can be a useful rehabilitation tool for individuals with elevated thresholds for vestibular motion perception, such as people with vestibulopathy (Eder et al. 2022), vestibular hypofunction (Priesol et al. 2014) or the elderly (Bermudez et al. 2016). In this respect, it remains to be seen whether noise such as that we applied here can improve not just vestibular perception, but also vestibulo-spinal function for balance control. This has been shown to be the case for GVS (Mulavara et al. 2011).

## Glossary

3D: three-dimensional
BRGLM: bias-reduced generalized linear model
95% CI: confidence interval at 95%
GLM: general linear model
GLMM: Generalized Linear Mixed Model
GVS: galvanic vestibular stimulation
LIA: lapse-identification algorithm
RM-ANOVA: Repeated measures analysis of variance
SD: standard deviation
SR: stochastic resonance

## ACKNOWLEDGMENTS

We thank Claudia Brunetti, Giorgio Capuzzi, Alessia Celli and Greta Dimasi for help with the setup and experiments. This work was supported by the Italian Ministry of Health (Ricerca corrente, IRCCS Fondazione Santa Lucia, Ricerca Finalizzata RF-2018-12365985), Italian Space Agency (grant I/006/06/0 and grant 2019-11-U.0), INAIL (BRIC 2019), and Italian University Ministry (PRIN grant 20208RB4N9_002 and 2020EM9A8X_003).

## AUTHORS CONTRIBUTIONS

All authors designed the experiments, analyzed the results, and wrote the paper.

